# Asthma and affective traits in adults: a genetically informative study

**DOI:** 10.1101/530865

**Authors:** Kelli Lehto, Nancy L. Pedersen, Catarina Almqvist, Yi Lu, Bronwyn K Brew

## Abstract

Depression, anxiety and high neuroticism (affective traits) are often comorbid with asthma. A causal direction between the affective traits and asthma is difficult to determine, however, it may be that there is a common underlying pathway attributable to shared genetic factors. Our aim was to determine whether a common genetic susceptibility exists for asthma and each of the affective traits.

An adult twin cohort from the Swedish Twin Register underwent questionnaire-based health assessments (n=23 693) and genotyping (n=15 908). Firstly, questionnaire-based associations between asthma and affective traits were explored. This was followed by genetic analyses: a) polygenic risk scores (PRS) for affective traits were used as predictors of asthma, and b) linkage-disequilibrium score regression based on genome-wide association results from UK Biobank was used to quantify genetic correlations.

Analyses found that the questionnaire-based associations between asthma and each affective trait were associated (OR 1.7, 95%CI 1.5-1.9 major depression, OR 1.5, 95%CI 1.3-1.6 anxiety, and OR 1.6, 95% 1.4-1.8 high neuroticism). Genetic susceptibility for neuroticism explained the variance in asthma with a dose response effect; that is, those in the highest neuroticism PRS quartile were more likely to have asthma than those in the lowest quartile (OR 1.4, 95%CI 1.2-1.6). Genetic correlations were found between depression and asthma (r_g_= 0.17), but not for anxiety or neuroticism score.

We conclude that the observed comorbidity between asthma and the affective traits may in part be due to shared genetic influences between asthma and depression and neuroticism, but not anxiety.

## INTRODUCTION

Asthma is a complex disease involving inflammation of the lungs and airways. It is the most common chronic respiratory disease in adults worldwide with a prevalence of 4-8% [1, 2]. Adults with asthma have twice as high risk of also having depression or anxiety compared to the general population [3-7]. This can lead to lowered quality of life and worse health outcomes. In particular, having depression can lead to poor adherence to inhaled corticosteroids, poor asthma control, and increased asthma exacerbations [8-10]. Despite a number of studies investigating the link between asthma, depression and anxiety, direction of causality has been difficult to determine [11], hence leading to the conclusion that perhaps a common underlying pathway exists [12].

Similarly, several studies have shown that the personality trait neuroticism influences the risk for asthma in adults, possibly increasing the risk up to three times among those with high neuroticism scores [13-15]. Neuroticism is characterized by a tendency to experience emotional instability and excessive worry, and has notable stability across the lifespan [16, 17]. It is also often a predecessor of depression and anxiety, which are considered to be psychiatric diagnoses that can change throughout a person’s lifetime. All three affective traits show considerable correlations on both phenotypic and genotypic levels [18, 19].

A common genetic pathway seems to be a plausible explanation for co-occurrence between asthma and the affective traits especially as adult asthma is moderately heritable, as are depression, anxiety and neuroticism [20-22]. A Finnish study of adult twins found some evidence for a genetic association between asthma and anxiety/depression [23]. Our own familial aggregation study in twin children suggested a common underlying cause for asthma and anxiety/depression, however a genetic hypothesis was not supported [24]. Both of these studies may have been underpowered and relied on twin modelling based on within-pair comparisons rather than using genotypic information. Furthermore, a shared genetic susceptibility between asthma and neuroticism has not been investigated at molecular genetic level to date. More direct genetic research is needed to assess whether there is genetic variance in common to affective traits and asthma using recent information from powerful genome-wide association studies (GWASs) for each of these conditions.

The objectives of this study are to: 1) confirm associations between asthma and affective traits (depression, anxiety and neuroticism) and 2) investigate whether there is evidence for a shared genetic relationship between each of the affective traits and asthma in adults. This will be done by using results from large GWASs to investigate cross-trait genetic association using a polygenic risk score approach and to estimate genetic correlations between traits using linkage disequilibrium score regression (LDSC).

## METHODS

### Study Population

Data were derived from three data collection occasions in the Swedish Twin Register [25-27] (Figure 1). In 1972-1973 *like*-sexed twins born 1926-58 were mailed a questionnaire collecting demographic, medical, pollution exposure, life-style and personality information including about neuroticism. About 36 000 twins responded (aged 15-47) [25]. These were followed up in 1998-2002 (ages 41-74 years) with a computer assisted telephone interview including questions regarding common complex diseases such as asthma, depression and anxiety. There was a 74% response rate of those alive and living in Sweden, n= 23 693 [26]. In addition, at this time *opposite* sex pairs were recruited for the telephone interview, increasing the number to a total of 38 633 participants. Between 2002-2010 (ages 44 to 82) these same twins that were still alive and had not participated in other genetic studies were asked to donate blood for bio-banking and genotyping purposes [27]. Approximately half of those contacted gave blood samples [27] Genotype information is now available for 15 908 of these.

**Figure 1.**
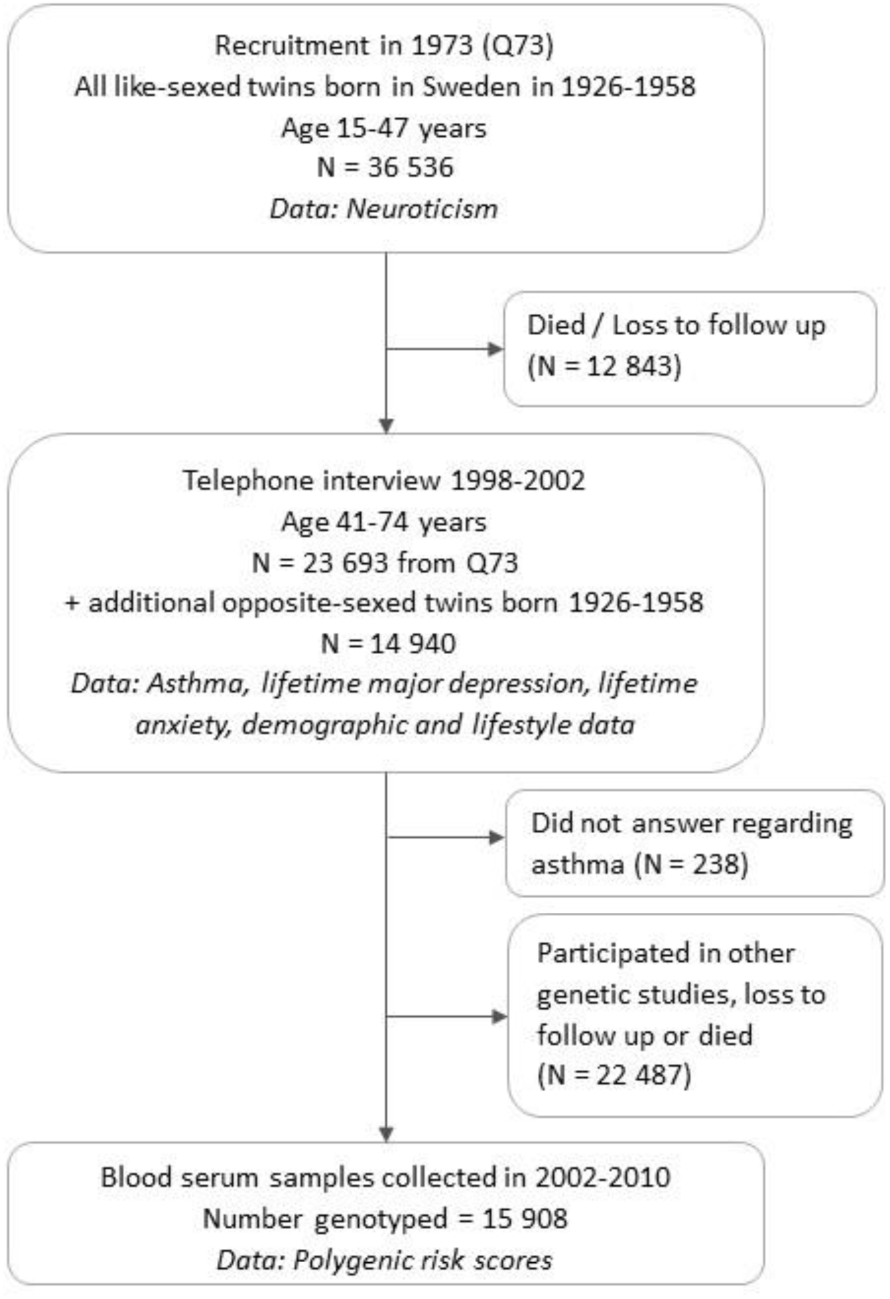
Flowchart of participation in the Swedish Twin register, birth years 1926-1958

## Measures

### Asthma

Asthma is a derived variable based on a number of asthma and allergy questions reflecting asthma ever in their lifetime. We scored positive for asthma those that either reported that their *asthma had been diagnosed by a doctor* OR they had *probable asthma based*. To be positive for *probable asthma* the subject had to have **one** of: a) self-reported asthma; whistle breathing; breathlessness at rest or breathlessness on waking up, AND **one** of b) asthma triggered by pollen, fur or hay; asthma onset between 4 and 30 years; self-reported hayfever with a debut between 4-30 years; asthma onset < 4 years and still have asthma/ age last had asthma was 4-20 years [28].

### Affective traits

Major Depression is a derived variable based on a number of questions based on the Composite International Diagnostic Interview Short Form (CIDI-SF). Positive for Major Depression is given if there are: symptoms of major depression (dysphoria and anhedonia) AND a CIDI-SF sum score ≥ 4 or positive for antidepressant use [29].

Self-report anxiety was a positive answer to the question ‘Have you ever had a period lasting one month or longer when most of the time you felt worried and anxious?’.

A short form of the Eysenck Personality Questionnaire was used to measure neuroticism [30]. Neuroticism was used both as a continuous trait and as a categorical trait divided into approximate quartiles (scores: 0-1, 2, 3-4, >5).

### Polygenic Risk Scores (PRS)

Single nucleotide polymorphism (SNP)-based genotyping was carried out using DNA from whole blood. Summary statistics from large GWAS (discovery sample) were used to compute genetic propensity scores for each trait in our own study population (target sample) using the individual genotypes. We calculated polygenic risk scores for major depressive disorder (PRS_MDD_), primary anxiety disorders (PRS_ANX_) and neuroticism (PRS_N_) by summing risk alleles (0, 1 or 2 alleles) at each SNP across the genome, weighted by individual SNP effect size derived from the discovery GWAS[18, 31, 32]. Polygenic risk scores were established at eight p-value thresholds (*p*T) ranging from 5 × 10^-8^ to 1 and then transformed to z-scores (see online data supplement for more details).

### Covariates

Age at interview, sex, years of education, body mass index (BMI) and current smoking were derived from the 1998-2002 questionnaire.

## Statistical Analysis

### Questionnaire-based associations

Multiple logistic regression models were used to determine the associations between each of the affective traits and asthma in the 23 693 twins with data available on all traits of interest. Models included: (1) unadjusted; (2) adjusted for potential confounders-age, sex and education; (3) adjusted for potential lifestyle mediators; (4) adjusted for neuroticism; and, (5) mutually adjusted for all affective traits to see if any one trait was driving the observed associations. We also stratified the analysis by sex. Relatedness of twin pairs was accounted for by including a standard error sandwich estimator based on twin pair ID.

### Polygenic Risk Score Analysis

First, the PRS’s for affective traits in our sample were validated (see online supplement for details). Then the association between questionnaire-based asthma and the polygenic risk score for each affective trait was tested in separate logistic regression models, adjusting for age, sex, 10 genetic ancestry principle components (PCs), genotyping array and relatedness. PCs are used to account for population stratification within the study population. The variance in asthma explained by each PRS was determined by comparison of Nagelkerke R^2^ from full (PRS and covariates) and reduced (covariates only) models. False discovery rate (FDR) at < 0.05 was applied to deal with multiple testing over eight p-value thresholds. Next, the PRS *p*_T_ explaining the greatest asthma variance was divided into quartiles based on distribution of PRS *p*_T_. Odds ratios were computed comparing the first quartile (i.e. the lowest genetic risk for affective traits) to other three quartiles in logistic regression models with asthma as an outcome, adjusted as above.

### Genetic correlations

In a separate analysis based on GWAS summary statistics from the UK biobank, genetic correlations between asthma and affective traits were estimated using LDSC without needing individual-level genotype information. Association test statistics are regressed on their LD scores, a measure of each SNP’s relationship with other variants [33]. The analyses were conducted in the LD Hub environment [34]. The most recently published GWAS summary statistics for asthma were used [35], restricted to samples with European descent. GWAS summary statistics for the affective traits were based on UK Biobank data available from http://www.nealelab.is/uk-biobank/. These included: two composite questions regarding a doctor or psychiatrist visit for anxiety/ depression/ tension/ nerves; self-reported depression; self-reported anxiety/panic attacks and neuroticism score. Because neuroticism score is constructed from items tapping into depression symptoms and worry /anxiety [36], we also analyzed all 12 neuroticism items to test which items show the highest genetic correlations with asthma. See the online data supplement Table E1 for a more detailed list of variables and GWAS sample sizes.

### Sensitivity Analyses

Associations were tested for *probable asthma* to check that the associations did not differ from our derived asthma variable that included asthma diagnoses, in case COPD cases had been misdiagnosed as asthma.

Data management was conducted using SAS 9.4. Statistical analysis was conducted using STATA/IC 15.1.

The study was approved the Regional Ethical Review Board in Stockholm, Sweden.

## RESULTS

The population of twins used for the questionnaire-based association analyses had a mean age of 56.0 years (SD 7.92 years) at testing in 1998-2002. The prevalence of asthma in this population was 7.9% (n=2137), and 40% of these reported both a diagnosis and symptoms (probable asthma), 33% only a diagnosis and 27% only symptoms. The prevalence of major depression was 20.1% and anxiety 19.9%. The descriptive characteristics and affective traits for those with and without asthma is shown in Table 1.

**Table 1.** Descriptive features of the study populations.

|  |  | Phenotypic analysis population |  | Genetic analysis (PRS) population |  |
| --- | --- | --- | --- | --- | --- |
|  |  | Asthma<br>N = 2,135 | No Asthma<br>N = 21,414 | Asthma<br>N = 1,448 | No Asthma<br>N = 14,454 |
| Age | Mean (SD) | 55.03 (7.98) | 56.11 (7.94) | 53.57 (7.13) | 54.93 (7.34) |
|  | Range | 41 - 74 | 41 - 74 | 41 - 74 | 41 - 74 |
| Sex – n (%) |  |  |  |  |  |
|  | Males | 913 (42.76%) | 10,167 (47.48%) | 600 (41.44%) | 6,976 (48.26%) |
|  | Females | 1,222 (57.24%) | 11,247 (52.52%) | 848 (58.56%) | 7,478 (51.74%) |
| Level of Education - n (%) |  |  |  |  |  |
|  | > 9 yrs | 567 (26.56%) | 5,357 (25.02%) | 460 (31.77%) | 4,274 (29.57%) |
|  | ≤ 9 yrs | 1,417 (66.37%) | 14,809 (69.16%) | 878 (60.64%) | 9,360 (64.76%) |
|  | Missing | 151 (7.07%) | 1,248 (5.83%) | 110 (7.60%) | 820 (5.67%) |
| Smoking - n (%) |  |  |  |  |  |
|  | Yes | 410 (19.2%) | 4,395 (20.52%) | 228 (15.75%) | 2,582 (17.86%) |
|  | No | 1,719 (80.52%) | 16,951 (79.16%) | 1,220 (84.25%) | 11,862 (82.07%) |
|  | Missing | 6 (0.28%) | 68 (0.32%) | 0 | 10 (0.07%) |
| BMI | N | 2,103 | 21,041 | 1,437 | 14,335 |
|  | Mean (SD) | 25.81 (4.081) | 24.94 (3.46) | 25.55 (3.72) | 24.87 (3.33) |
|  | Range | 14.7 – 47.8 | 14.0 – 47.8 | 15.0 - 45.5 | 16.2 - 47.3 |
| Major Depression – n (%) |  |  |  |  |  |
|  | Yes | 629 (29.46%) | 4,103 (19.16%) | 450 (31.08%) | 2,999 (20.75%) |
|  | No | 1,429 (66.93%) | 16,537 (77.23%) | 980 (67.68%) | 11,292 (78.12%) |
|  | Missing | 77 (3.61%) | 774 (3.61%) | 18 (1.24%) | 163 (1.13%) |
| Anxiety – n (%) |  |  |  |  |  |
|  | Yes | 562 (26.32%) | 4,103 (19.16%) | 402 (27.76%) | 2,865 (19.82%) |
|  | No | 1,541 (72.18%) | 17,103 (79.87%) | 1,037 (71.62%) | 11,486 (79.47%) |
|  | Missing | 32 (1.50%) | 208 (0.97%) | 9 (0.62%) | 103 (0.71%) |
| Neuroticism | N | 1,902 | 19,181 | 811 | 8,463 |
|  | Mean (SD) | 3.26 (2.49) | 2.73 (2.30) | 3.20 (2.43) | 2.70 (2.27) |
|  | Range | 0 - 9 | 0 - 9 | 0 - 9 | 0 - 9 |
|  | Quartile 1 | 575 (26.93%) | 7,177 (33.52%) | 241 (16.64%) | 3,192 (22.08%) |
|  | Quartile 2 | 294 (13.77%) | 3,136 (14.64%) | 136 (9.39%) | 1,373 (9.50%) |
|  | Quartile 3 | 458 (21.45%) | 4,714 (22.01%) | 201 (13.88%) | 2,108 (14.58%) |
|  | Quartile 4 | 575 (26.93%) | 4,154 (19.40%) | 233 (16.09%) | 1,790 (12.38%) |
|  | Missing | 233 (10.91%) | 2,233 (10.43%) | 637 (43.99%) | 5,991 (41.45%) |

The population of twins used for the asthma – polygenic risk score association tests had a mean age of 54.8 years (SD 7.37) at testing in 1998-2002. The descriptive characteristics and prevalences were similar to the population used for the questionnaire-based analyses (Table 1). The prevalence of asthma in this population was 9.0%, major depression 21.7% and anxiety 20.5%.

Questionnaire-based associations between each of major depression, anxiety, neuroticism (both continuous and categorical) and asthma are reported in Table 2. The associations remained significant after adjusting for potential confounders (Model 2) and mediators (Model 3). Associations were slightly stronger for men than for women (Table 2). When the models for major depression and anxiety were also adjusted for neuroticism the associations slightly reduced but the significance stayed the same (Table 2, Model 4). Mutual adjustment for all three affective traits seemed to attenuate the association between anxiety and asthma (except in males) and only slightly lowered the associations for major depression and neuroticism (continuous) (Table 2, Model 5).

**Table 2.**
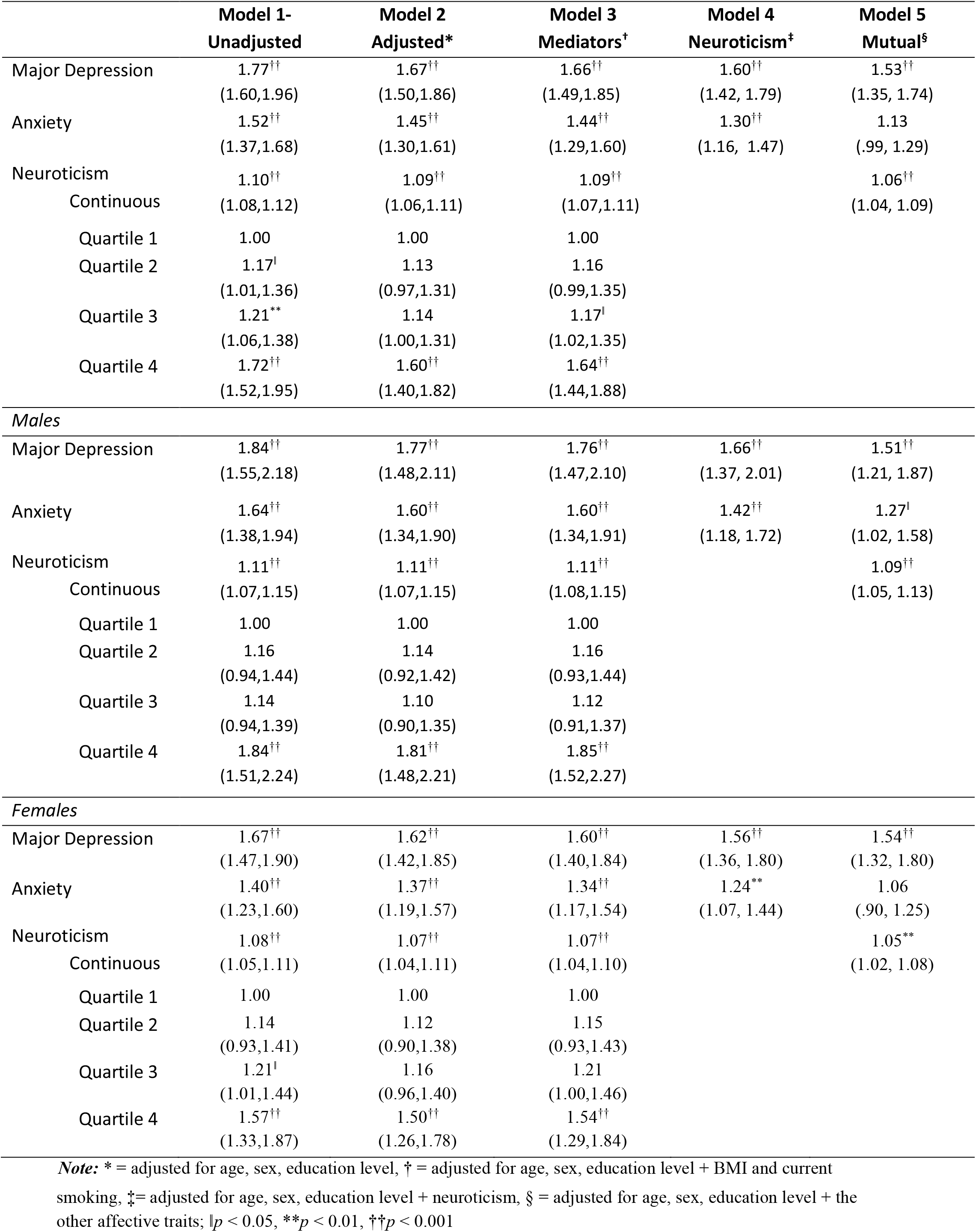
Questionnaire-based associations between affective traits and asthma for the whole population and separately for men and women. (OR and 95% CI)

The sensitivity analysis using the probable asthma definition did not change the effect sizes (see Table S2 in the online data supplement).

Validation of polygenic risk scores for major depression, anxiety and neuroticism confirmed them as significant predictors of each respective trait in our study population (See Figure E1). The PRS_N_ explained up to 0.2% of the variance in asthma (p<0.001), which was about half of the variance explained by the genetic score in neuroticism itself (Figure 2). No associations were seen for PRS_MDD_ and PRS_ANX_ with asthma. One *p*_T_ of each three affective trait was selected for subsequent analysis in quartiles, based on the highest variance explained in asthma, which was *p*_T_ 0.5 for all three traits.

**Figure 2.**
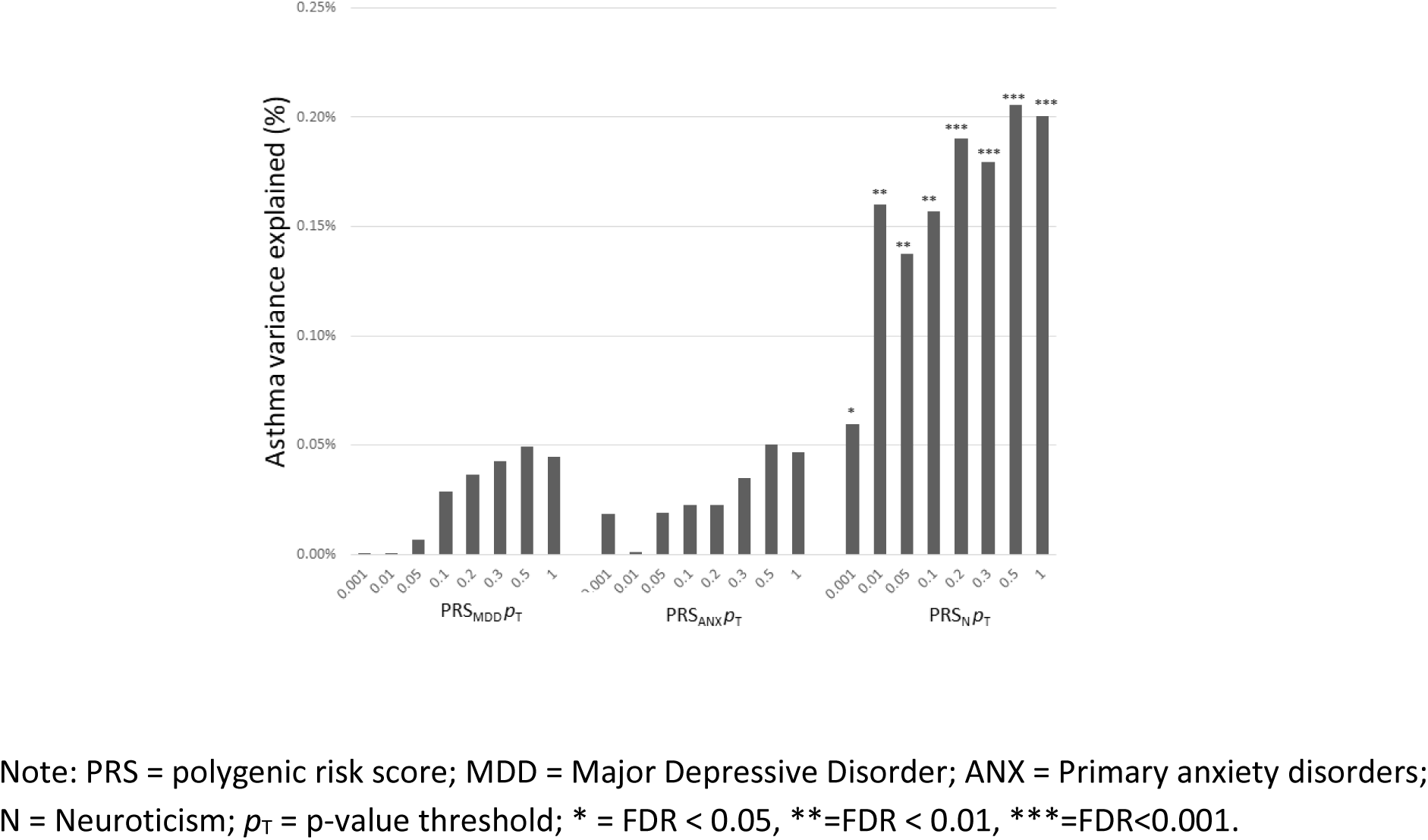
Polygenic risk scores for each of the affective traits explaining variance in asthma (n = 15,908) Note: PRS = polygenic risk score; MDD = Major Depressive Disorder; ANX = Primary anxiety disorders; N = Neuroticism; *p*_T_ = p-value threshold; * = FDR < 0.05, **=FDR < 0.01, ***=FDR<0.001.

Quartile-based analysis revealed a dose-response effect for neuroticism risk scores, that is, with each increasing quartile of neuroticism score the risk of asthma increased (Table 3). For example, individuals with the highest neuroticism scores in the top quartile had a 37% higher odds (p<0.001) of having asthma compared to the group with the lowest genetic risk. No elevated asthma risk was observed with increasing genetic predisposition for MDD or anxiety.

**Table 3.** Associations of affective traits’ PRS quartiles (compared to the lowest genetic risk quartile) for each of the affective traits and asthma. (Odds ratios and 95% confidence intervals)

|  | PRS*<br>quartile | OR | 95% CI |  | p-value |
| --- | --- | --- | --- | --- | --- |
|  |  |  | Lower | Upper |  |
| Major Depression | Quartile 2 | 0.94 | 0.80 | 1.10 | 0.45 |
|  | Quartile 3 | 1.04 | 0.88 | 1.21 | 0.67 |
|  | Quartile 4 | 1.12 | 0.96 | 1.31 | 0.17 |
| Anxiety | Quartile 2 | 1.12 | 0.96 | 1.32 | 0.16 |
|  | Quartile 3 | 1.14 | 0.97 | 1.34 | 0.11 |
|  | Quartile 4 | 1.12 | 0.96 | 1.32 | 0.16 |
| Neuroticism | Quartile 2 | **1.18** | **1.00** | **1.39** | **0.05** |
|  | Quartile 3 | **1.25** | **1.06** | **1.48** | **0.01** |
|  | Quartile 4 | **1.37** | **1.17** | **1.61** | **<0.001** |

**Note:** \* = The threshold probabilities selected were $p_T$ 0.5 for all three traits.

The LD score regression found that asthma was genetically correlated with the composite questions regarding a visit to a doctor or psychiatrist for anxiety, depression, tension or nerves (Table 4). There was also a modest genetic correlation of asthma with self-reported depression (r_g_ = 0.17), but not with anxiety or neuroticism score. Item-level analysis for neuroticism revealed genetic correlation between asthma and items tapping into depressed mood (fed-up feelings, loneliness) rather than worry/anxiety.

**Table 4.**
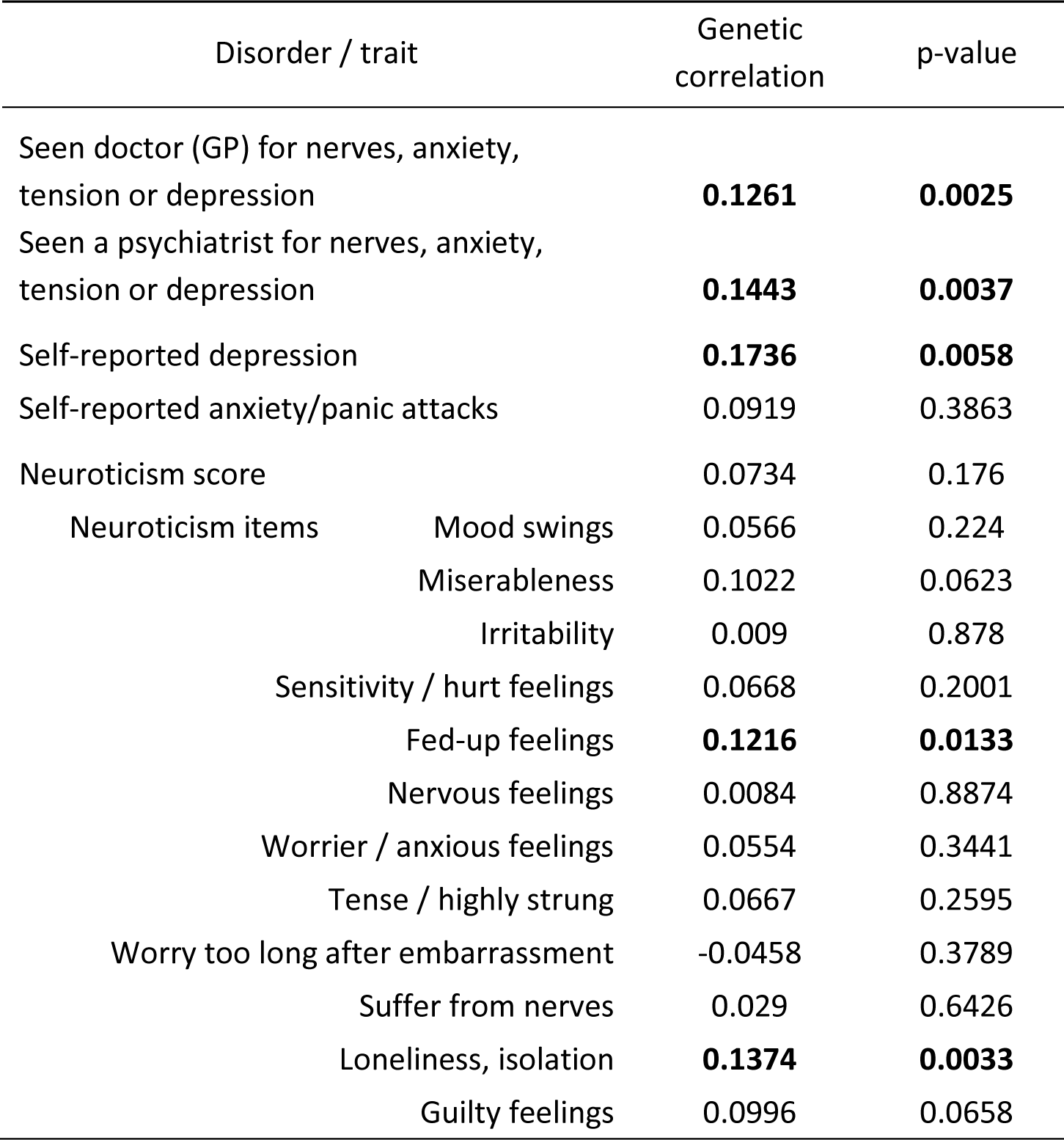
LD Score regression results between affective traits from UK Biobank and asthma from the Multiancestry Consortia association study [35]

## DISCUSSION

In this study we attempted to triangulate the role of genetic influences on the associations between asthma and affective traits by first investigating phenotypic associations and then applying two different genetic methods (polygenic risk score approach and LDSC) to investigate shared genetic relationship between asthma and affective traits. To the best of our knowledge, this is the first time a genetic link between asthma and depression, anxiety and neuroticism has been investigated using genome-wide data. The phenotypic results found associations between asthma and the affective traits-major depression, anxiety and neuroticism. The polygenic risk score analysis using genotypes from a large twin population found modest associations between asthma and genetic susceptibility for neuroticism. The LDSC using GWAS summary statistics from UK Biobank detected genetic overlap between asthma and depression.

We found higher asthma risk in individuals with major depression, which is in line with previous research [4, 5, 7]. Modest but significant genetic correlation between self-reported depression and asthma was found using the LDSC approach, suggesting possible genetic overlap and shared biological pathways between these disorders. Potential explanatory biological candidates for depression and asthma comorbidity have been previously reviewed by Jiang et al. and include specific cytokine inflammatory factors such as IL-1, IL-4, IL-7 and TNF-α [37]. However, we did not detect a significant association between asthma and polygenic risk score for major depressive disorder. The discordant results of the two methods could be potentially explained by different diagnostic criteria used in the two depression GWASs (diagnostic interview in PRS summary statistics *vs*. self-reported depression in UK Biobank) or differences in their power to detect genetic signals, UK Biobank having a larger sample size. The ability of polygenic risk scores to detect associations increases with increasing power of the discovery GWAS, therefore future replications are warranted with genome-wide association studies conducted in larger samples.

Anxiety has repeatedly been associated with asthma at the phenotypic level [3, 38, 39]. Our phenotypic results confirmed these associations in models where the effect of each affective trait was explored separately. However, considering the substantial correlation among anxiety, depression and neuroticism, we also conducted mutual adjustment for all affective traits. These results indicated a significant drop in the effect size of anxiety, but not in the other affective traits. This could potentially indicate that depression and neuroticism may have a stronger influence on asthma compared to anxiety. Similar patterns were also seen in both analyses using genetic data. First, polygenic risk scores for anxiety failed to predict asthma in our sample. These results could potentially be explained by the lower power of the anxiety PRS compared to the neuroticism PRS, due to the smaller sample size of the original GWAS for anxiety. However, the genetic correlation method, LDSC, did not suffer from these limitations since data were derived from the well-powered UK Biobank sample. Yet the results suggested the same pattern - significant genetic correlations were found between asthma and self-reported depression as well as depression-related neuroticism items, but no genetic correlation was found with self-reported anxiety nor anxiety-related neuroticism items. Nevertheless, these results should be replicated in the future, when more powerful genome-wide association studies on anxiety disorders emerge.

With regards to neuroticism, other studies have found up to a fifteen-year longitudinal association between neuroticism and the development of asthma [5, 13, 15]; our study extends this period up to 29 years. Even after this extremely long follow-up time, the association between neuroticism and asthma was still rather strong, with effects similar to concurrently measured major depression (lowest vs highest quartile). Neuroticism is known to be a relatively stable personality trait [16, 17] and therefore may also be associated with asthma throughout the life span. We were able to demonstrate that polygenic risk scores for neuroticism were associated with asthma, which may provide some explanation for such long-term associations. The effects of such genetic composite scores are small, but the predictive ability of polygenic risk scores will improve with increasing power of GWASs to detect very small genetic signals. However, in the LDSC analysis we did not find evidence for genetic overlap between neuroticism and asthma despite data from a larger resource compared to the GWAS used for creating the neuroticism polygenic risk score. The neuroticism score as measured in UK Biobank and used in this study for LDSC, is composed of 12 rather heterogeneous items, some capturing more depressed mood and others nervousness and anxiety. Recently it was reported that the genetic signal in neuroticism may originate from two different genetic clusters: depressed mood (‘mood swings, ‘loneliness’, ‘fed-up feelings’, ‘miserableness’) and worry (‘nervous feelings’, ‘worrier/anxious feelings’, ‘tense/highly strung’, ‘suffer from nerves’) [36, 40]. Therefore, we investigated the genetic correlations for each neuroticism item separately and indeed found depression-related items to be more strongly associated with asthma compared to worry / anxiety related items. This is further suggests that the genetic association may be stronger between asthma and depression than with anxiety.

This study has some limitations. The weak associations of anxiety in comparison to the other two affective traits could be influenced by the single-item anxiety measure that was used in the questionnaire-based association testing. The assessment of major depression and neuroticism were based on composite questions and therefore might be more robust. However, the LDSC analysis, which did not suffer from this limitation, still suggested the same pattern. Another limitation is that we were not able to test the direction of the association between asthma and affective traits, despite the longitudinal association found between neuroticism and asthma.

### Conclusion and Future Directions

In conclusion, the observed comorbidity between asthma and the affective traits is in part due to genetic influences on the affective traits, and a moderate overlap in genes of importance for both asthma and depression and neuroticism but not anxiety. This is the first study to observe a genetic susceptibility between asthma and affective traits. Replication studies will be advantageous in the future using more powerful GWAS once they become available. Furthermore, this study provides endorsement for future basic science research into the specific shared biological pathways for inflammatory and psychiatric disorders.

## Acknowledgements

We acknowledge The Swedish Twin Registry at the Karolinska Insitutet for access to data and the database managers who have worked on the registry over several decades. We also acknowledge all participants in the Swedish Twin Register without whom this study would not be possible. We thank Robert Karlsson at the Karolinska Institutet for biostatistical advice.

