## Supplementary info for "Asthma and affective traits in adults: a genetically informative study"

### Online Data Supplement.

### Supplementary Methods

**Validation of PRS Scores**

To validate PRS's for affective traits in our sample, linear (Neuroticism) or logistic (MDD and anxiety) regression was run on each PRS $p_T$ of each trait on respective phenotypes (i.e. $PRS_{MDD}$ predicting lifetime major depression in our population) (Figure E1). Regressions were adjusted for age, sex, 10 genetic ancestry Principal Components (PCs), genotyping array and relatedness. PCs are used to account for population stratification within the study population.

**Polygenic Risk Scores**

Whole blood collected 2002-2010 was used for genotyping on two genotyping platforms: the Illumina OmniExpress bead chip, further imputed to Hapmap 2 build 36 reference panel and the Illumina Infinium PsychArray-24 BeadChip, further imputed to 1000 Genomes Phase 3 reference panel (1). For each of the MZ pairs only one twin was genotyped and the genotype information was imputed for the co-twin.
The PRS for neuroticism ($PRS_N$), major depression disorder ($PRS_{MDD}$) and primary anxiety disorders ($PRS_{ANX}$) in this study were created using available summary statistics with all Swedish samples excluded to avoid potential sample overlap between the discovery and the target samples. Such sample overlap may lead to significant inflation in effect sizes. The sample sizes for the GWAS meta-analyses were as follows: MDD = 38 695 (PGC only), primary anxiety disorders = 17 310, neuroticism = 166 005. In order to deal with linkage disequilibrium, clumping was performed on the discovery association data. This procedure selects most significantly associated SNPs and excludes SNPs in strong LD ($r2 < 0.1$ within a 1000 kb window using Plink version 1.9). The maximum number of LD-pruned SNPs considered ranged from 84,298 to 112,719, depending on the affective trait and genotyping platform.

**UK Biobank variables**

A selection of variables from the UK Biobank, capturing neuroticism and ever having problems with depression or anxiety, were used in LDSC analysis. The items and each respective item number in UK Biobank are reported in Table E1.

### Supplementary results

**Table E1.** UK Biobank variables used in LDSC analysis.

| Trait | Item number in UK Biobank | GWAS sample size |
|---|---|---|
| Neuroticism score | 20127 | 274 108 |
| Neuroticism items: | | |
| Does your mood often go up and down? | 1920 | 329 428 |
| Do you ever feel 'just miserable' for no reason? | 1930 | 331 856 |
| Are you an irritable person? | 1940 | 322 668 |
| Are your feelings easily hurt? | 1950 | 327 832 |
| Do you often feel 'fed-up'? | 1960 | 330 549 |
| Would you call yourself a nervous person? | 1970 | 328 725 |
| Are you a worrier? | 1980 | 328 717 |
| Would you call yourself tense or 'highly strung'? | 1990 | 327 232 |
| Do you worry too long after an embarrassing experience? | 2000 | 323 766 |
| Do you suffer from 'nerves'? | 2010 | 325 248 |
| Do you often feel lonely? | 2020 | 332 263 |
| Are you often troubled by feelings of guilt? | 2030 | 328 769 |
| Non-cancer illness code_ self-reported: depression | 20002_1286 | 337 159 |
| Non-cancer illness code_ self-reported: anxiety/panic attacks | 20002_1287 | 337 159 |
| Seen doctor (GP) for nerves_ anxiety_ tension or depression | 2090 | 335 108 |
| Seen a psychiatrist for nerves_ anxiety_ tension or depression | 2100 | 335 888 |

**Table E2.** Phenotypic Associations between affective traits and probable asthma. (OR and 95% CI)

| | **Model 1- unadjusted** | **Model 2- confounders** adjusted for age, sex, SES | **Model 3- mediators** adjusted for age, sex, SES, BMI, smoking |
|---|---|---|---|
| Neuroticism | $1.092^{***}$ (1.067,1.117) | $1.080^{***}$ (1.054,1.107) | $1.083^{***}$ (1.056,1.110) |
| Anxiety | $1.521^{***}$ (1.353,1.709) | $1.431^{***}$ (1.264,1.619) | $1.425^{***}$ (1.258,1.615) |
| Major Depression | $1.762^{***}$ (1.568,1.980) | $1.618^{***}$ (1.429,1.832) | $1.598^{***}$ (1.410,1.812) |
| *Males* | | | |
| Neuroticism | $1.104^{***}$ (1.065,1.144) | $1.099^{***}$ (1.058,1.140) | $1.100^{***}$ (1.059,1.142) |
| Anxiety | $1.806^{***}$ (1.505,2.168) | $1.646^{***}$ (1.364,1.987) | $1.646^{***}$ (1.363,1.988) |
| Major Depression | $1.914^{***}$ (1.587,2.307) | $1.721^{***}$ (1.416,2.090) | $1.695^{***}$ (1.394,2.061) |
| *Females* | | | |
| Neuroticism | $1.086^{***}$ (1.053,1.119) | $1.070^{***}$ (1.036,1.105) | $1.073^{***}$ (1.039,1.108) |
| Anxiety | $1.359^{***}$ (1.166,1.584) | $1.298^{**}$ (1.105,1.524) | $1.289^{**}$ (1.095,1.517) |
| Major Depression | $1.692^{***}$ (1.452,1.971) | $1.557^{***}$ (1.328,1.826) | $1.538^{***}$ (1.309,1.807) |

$^{*}$ $p < 0.05$, $^{**}$ $p < 0.01$, $^{***}$ $p < 0.001$

**Figure E1.** PRS for affective traits explaining variance in their respective phenotypes (Neuroticism, Anxiety and Major Depression) in the target sample, n = 10,075

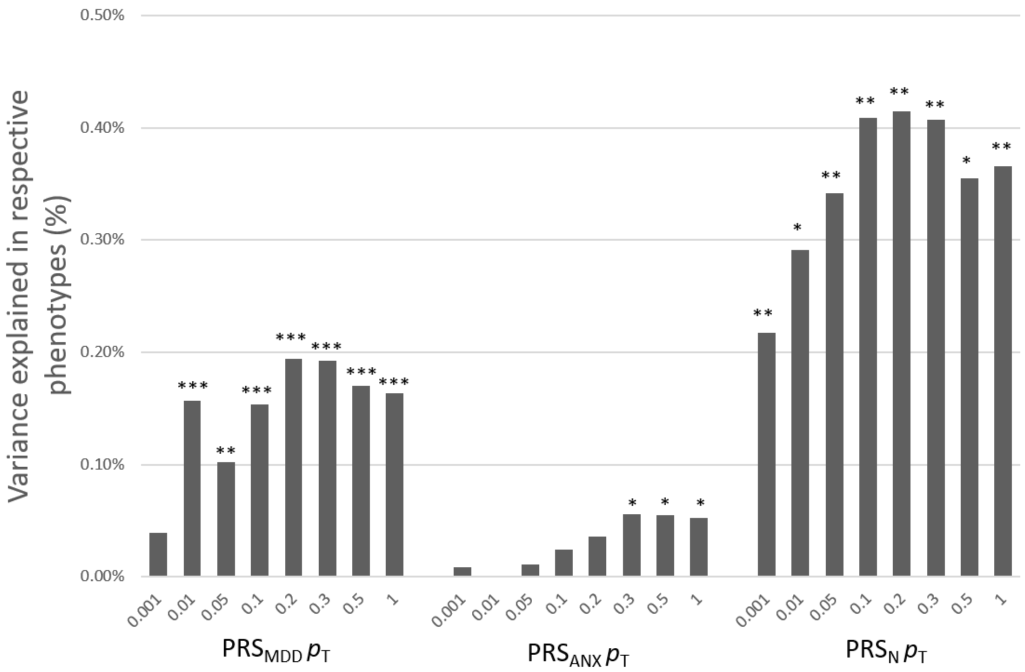


*Note:* PRS = polygenic risk score; MDD = Major Depressive Disorder; ANX = Primary anxiety disorders; N = Neuroticism; $p_T$ = p-value threshold; Uncorrected p-values; $^{*}p < 0.05$, $^{**}p < 0.01$, $^{***}p < 0.001$.
